# Molecular Plasticity of T Cells Informs Their Possible Adaptation in 4T1 Tumors

**DOI:** 10.1101/2025.10.17.683070

**Authors:** Md. Iftehimul, Robert H. Newman, Scott H. Harrison, Roshonda B. Jones, Perpetua M. Muganda, Bryan L. Holloman, Muhammad T. Hossain, Checo J. Rorie, Misty D. Thomas, Joseph L. Graves, Howard L. Kaufman, Dipongkor Saha

**Author notes:** Correspondence to: Dipongkor Saha.

## Abstract

**Background:** The triple-negative breast cancer (TNBC) microenvironment (TME) undergoes progressive reprogramming, transitioning from an early immune-active state to a late immune-suppressed state. While tumor cell plasticity has been extensively studied, the molecular plasticity of T cells *in vivo* remains poorly defined.

**Objectives:** To characterize transcriptional changes in T cells during TNBC progression and identify stage-specific shifts in T cell function, polarization, and antigen-presenting cell (APC)-T cell interactions.

**Results:** Transcriptional analysis of T cells from BALB/c mice bearing 4T1 tumors at 1, 3, and 6 weeks revealed a decline in T cell-associated genes from 194 at 1 week to 156 at 6 weeks, with a significant late-stage loss of TCR diversity and contraction of natural killer T (NKT)- and γδ T cell-related transcripts. Cytokine and transcription factor dynamics reflected temporal T cell polarization: early (1 week) IL-12α/β-STAT4 signaling supports CD4^+^ type 1 T helper cell (Th1) and type 1 CD8^+^ cytotoxic T cell (Tc1) responses; intermediate (3 weeks) IL-21 and BCL6 expression suggest transient CD8^+^ cytotoxic follicular T cell (Tfc) skewing; and late (6 weeks) AhR and IL-1β induction reflect interleukin 17/22 producing CD8^+^ T cell (Tc17/Tc22) transition. Pro-inflammatory cytokines and chemokines increased over time, while immunosuppressive mediators (e.g., IL-10) declined significantly. Antigen-presenting cell (APC)-T cell crosstalk deteriorated at 6 weeks, characterized by a reduction in the expression of co-stimulatory and APC genes. Despite an early dominance of M1-like macrophage signals (e.g., IL-12α/β), persistent expression of arginase 1 (ARG1) and other M2-associated genes indicated a stable tolerogenic niche.

**Conclusions:** TNBC progression is characterized by progressive T cell functional decline, narrowing of TCR diversity, impaired APC-T cell interactions, and sustained macrophage-driven immunosuppression. These temporally coordinated immune shifts suggest tumor-driven adaptation toward immune evasion and identify potential windows for stage-specific immunotherapeutic intervention.

## Introduction

Triple-negative breast cancer (TNBC), defined by the absence of estrogen receptor (ER), progesterone receptor (PR), and human epidermal growth factor receptor 2 (HER2), represents one of the most clinically challenging subtypes of breast cancer^1^. This receptor-deficient profile renders TNBC unresponsive to endocrine or HER2-directed therapies. TNBC is particularly aggressive, with high recurrence rates and poor prognosis, especially in the metastatic setting^2^. Immunotherapies, such as immune checkpoint blockade, antibody-drug conjugates, and cancer vaccines, are emerging treatment options for TNBC. However, their efficacy relies heavily on the functional state of the host immune system, particularly tumor-infiltrating T lymphocytes (TILs) ^3,4^. Within the tumor microenvironment (TME), TILs encounter strong immunosuppressive cues that limit their ability to mount effective antitumor responses^5^.

T cells show a remarkable degree of molecular plasticity, undergoing dynamic and reversible changes in phenotype, function, gene expression, and metabolism in response to extrinsic stimuli^6,7^. This adaptability is fundamental for immune surveillance, enabling T cells to recognize diverse pathogens and operate across tissue contexts. In cancer, however, the TME hijacks T cell plasticity to promote immune evasion. For instance, cytotoxic CD8^+^ T cells often become dysfunctional or exhausted due to chronic antigen exposure, inhibitory ligands such as programmed death-ligand 1 (PD-L1), and nutrient depletion^8,9^. CD4^+^ helper T cells may be skewed toward regulatory T cells (Tregs) under the influence of transforming growth factor beta (TGF-β) and interleukin 10 (IL-10), thereby dampening immune surveillance^10,11^. T cells also shape tumor evolution. For instance, interferon-gamma (IFN-γ), traditionally viewed as an antitumor cytokine^12^, can paradoxically promote tumor cell survival, epithelial-to-mesenchymal transition (EMT), and metastasis^13-16^. This reciprocal relationship highlights that T cell plasticity is not only a reflection of immune dysfunction but also an engine of tumor adaptation. A better understanding of T cell plasticity in tumors could shed light on how tumors evade immune recognition and how immune cells might be reprogrammed to overcome these barriers. Therapies that limit harmful plasticity (e.g., preventing T cell exhaustion or Treg conversion) or promote beneficial reprogramming (e.g., enhancing memory T cell formation, metabolic fitness) are promising avenues to improve immunotherapy outcomes. While many studies have profiled tumor gene expression under drug pressure or other external interventions^17-22^; fewer have focused on untreated tumor progression *in vivo*. Previously, we investigated the molecular phenotypic plasticity of 4T1 TNBC cells, a well-characterized murine breast cancer model recapitulating many features of human TNBC, and their adaptation *in vivo*^23^. Here, we investigated the molecular phenotypic plasticity within the untreated 4T1 TME. Our analysis highlights how evolving T cell states can influence tumor adaptation, progression, and responsiveness to immunotherapies.

## Methods

To assess T cell plasticity in 4T1 bulk tumors over time *in vivo*, we retrieved paired-end RNA-seq sequencing data from nine samples belonging to the NCBI BioProject PRJNA588756, which included bulk tumor samples derived from the same batch of 4T1 cells implanted orthotopically into the mammary fat pad of BALB/c mice, collected at 1 week (early), 3 weeks (intermediate), and 6 weeks (late) post-tumor implantation^24^. We additionally used 4T1 cells, which are syngeneic to BALB/c mice, as a baseline in this study and these were retrieved from NCBI BioProjects PRJNA671832, PRJNA873199 and PRJEB36287. (**Table S1**). Transcriptomic sequence reads were quality-checked with FastQC v.0.12.1^25^, subsequently trimmed using Fastp v.0.24.0^26^ and aligned to the GRCm39 mouse reference genome (ensembl release 115) using HISAT2^27^. Gene-level read counts were quantified with FeatureCounts^28^. To test for differences in expression between baseline and each of the three time points, differential expression analysis was conducted in the R statistical program (v. 4.5.1) using the R package DESeq2^29^. After sequence count normalization, differentially expressed genes (DEGs) were identified based on a nominal p-value < 0.05 and an absolute log 2-fold change (Log2FC) ≥ 1. Upregulated and downregulated genes were classified using Log2FC ≥ 1 and Log2FC ≤ −1, respectively, with adjusted p-values also calculated. Default parameters were used for all aforementioned software. The log2FC values, unless otherwise specified, were indicated in parentheses in the Results and Conclusions sections. DEGs were filtered to only include those that were T cell-related genes. T cell-related genes were identified based on the nCounter Mouse PanCancer Immune Profiling Panel^30^ and cross-referenced with published literature to ensure biological relevance. The resulting DEGs were visualized in R using volcano plots and heatmaps^31^. For statistical comparisons, repeated-measures one-way analysis of variance (ANOVA) followed by Student-Newman-Keuls post hoc testing was performed using GraphPad Prism v10.5. Statistical significance was defined as *p ≤ 0.05, **p ≤ 0.01, ***p ≤ 0.001, and ****p ≤ 0.0001.

## Results

### Dynamic molecular plasticity of T cell profiles across stages of 4T1 tumor growth

Tumor plasticity not only reshapes cancer cells but also drives dynamic remodeling of the tumor immune microenvironment, particularly T cell compartments. To elucidate the plasticity of T cell compartments, we profiled key T cell-associated genes from 1 to 6 weeks of 4T1 tumor progression. Out of 200 T cell-specific genes outlined in the Mouse PanCancer Immune Profiling Panel^30^, 194 genes were expressed in 1-week tumors, decreasing modestly to 191 genes at 3 weeks and further to 156 genes at 6 weeks (**Fig. 1**). This gradual reduction suggests that T cell functionality and activity undergo dynamic modulation as tumors grow, reflecting a complex interplay between tumor growth and immune regulation^32^. The diminishing transcriptional breadth could be driven by impaired T cell receptor (TCR) signaling and shifts within specialized T cell subsets like Natural Killer T (NKT) and gamma delta (γδ) T cells^33^. To characterize the molecular plasticity of genes associated with NKT cell function, we profiled 26 key NKT-associated genes at 1-, 3-, and 6-weeks post-tumor implantation. Between 1 and 3 weeks, NKT cells demonstrated a robust effector phenotype, with high expression of *perforin 1* (*PRF1*; 12.44), *granzyme B* (*GZMB*; 13.06), *interferon-gamma* (*IFN-γ*; 6.88), *interleukin-13* (*IL-13*; 9.76), and *interleukin-21 receptor* (*IL-21R*; 11.90), indicating early regulatory and innate-like profiles. By 6 weeks, cytokine genes such as *IL-13* (5.69) were sharply downregulated, while IL-21 was lost, and cytotoxic expression of *PRF1* and *GZMB* persisted (**Table S2**). Statistical analysis revealed significant differences at 6 weeks compared to 1 and 3 weeks (p < 0.0001), whereas the comparison between 1 and 3 weeks was not significant (p = 0.2840) **(Fig. 2A)**. These results indicate that substantial molecular reprogramming of NKT cells occurs predominantly at later stages of tumor development. TCR-related variable (V) and joining (J) segment genes, including *TCR beta variable 2* (*TRβV2)* and *16 (TRβV16)*, and *TCR beta joining 1-1* (*TRβJ1-1)* and *2-1* (*TRβJ2-1*), became undetectable by 6 weeks, indicating narrowing TCR repertoire diversity (**Table S3**). No significant difference was observed between 1 and 3 weeks (p = 0.3159), suggesting stable TCR gene expression during the early to intermediate phases of tumor growth. Interestingly, at 6 weeks (p < 0.001; p < 0.0001), these genes were significantly decreased, suggesting a substantial reorganization of the TCR transcriptome by 6 weeks post-implantation. Conversely, *TCR alpha variable 1* (*TRαV1*) and *3-3* (*TRαV3-3*) emerged or persisted, suggesting a compensatory mechanism to sustain immune activity in a clonally restricted population (**Fig. 2B**).

**Figure 1.**
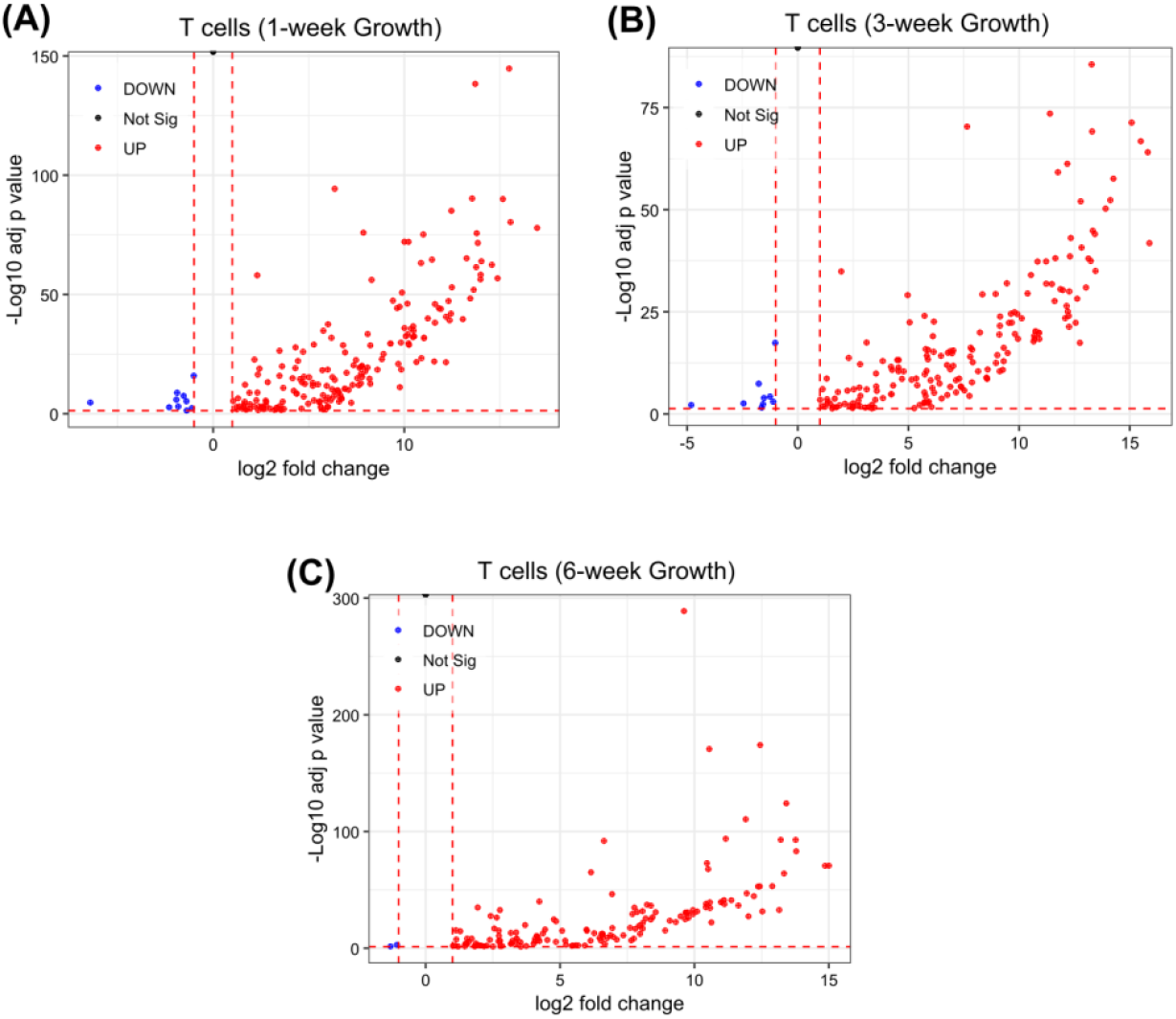
Volcano plots for DEGs that are T cell-associated genes, as determined by the Mouse PanCancer Immune Profiling Panel^30^, at different stages of 4T1 tumor growth. Compared to the 4T1 cell RNA-seq dataset as a baseline, these plots show gene expression changes at (A) 1 week (194 genes), (B) 3 weeks (191 genes), and (C) 6 weeks (156 genes) post-tumor implantation. The X-axis (log2FC) indicates the magnitude and direction of expression changes, with positive values denoting higher expression and negative values denoting lower expression, while the Y-axis (-Log10 adj p value) indicates the strength of the statistical significance of these expression changes.

**Figure 2.**
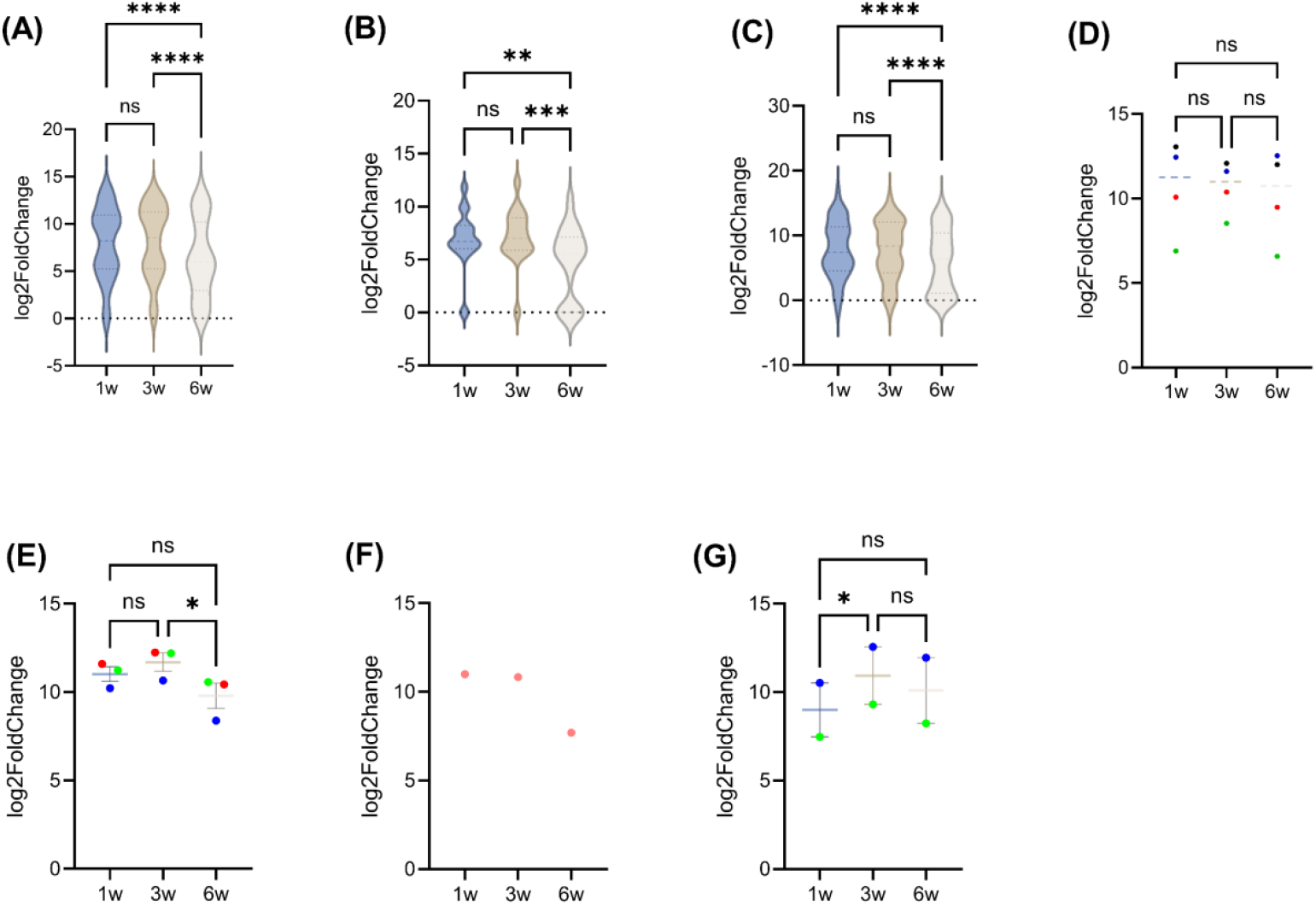
T cell expression profiles across different stages of tumor growth. Log2FC of various genes related to (A) natural killer T (NKT) cells, (B) T cell receptor (TCR), (C) gamma delta (γδ) T cells, (D) T cell activation markers (CD69 in red, IFNγ in green, PRF1 in blue, and GZMB in black), (E) CD3 isoforms (CD3ε in red, CD3γ in green, and CD3δ in blue), (F) CD4, and (G) CD8 isoforms (CD8α in blue and CD8β1 in green). ns, not significant; w, weeks; *p ≤ 0.05, **p ≤ 0.01, ***p ≤ 0.001, and ****p ≤ 0.0001. Figure A-C Genes presented in these figures are exactly refelcted in **Supplement tables S2-S4**.

The temporal plasticity of γδ T cells in the TME offers important insights for cancer immunotherapy^34^. In our study, γδ T cell-associated genes displayed significant transcriptional shifts at 6 weeks (p < 0.0001) compared to earlier time points. Several γδ TCR genes, such as *TCR gamma variable 1* (*TRγV1*), *TCR delta variable 4* (*TRδV4*), and *TCR delta variable 5* (*TRδV5*), and *SRY-box transcription factor 13* (*SOX13*) and *Zinc Finger and BTB Domain Containing 16* (*ZBTB16*), were downregulated or lost, indicating clonal contraction (**Figure 2C and Table S3**). However, cytotoxic mediators *PRF1* and *GZMB* remained relatively stable or slightly elevated, *IFN-γ* peaked transiently at 3 weeks (8.53) before declining at 6 weeks (6.58) **(Fig. 2D)**. T cell activation gene, such as *cluster of differentiation 69* (*CD69*), declined from 10.08 (earlier) to 10.38 (intermediate) to 9.49 (late stage). The *cluster of differentiation 3* isoforms (*CD3ε, CD3δ*, and *CD3γ*) also declined by late stage, indicating weakened TCR signaling **(Fig. 2E)**. *CD4* expression declined sharply at 6 weeks (∼11 to 7.69), and *CD8α*/*β1* peaked at 1-3 weeks before decreasing at 6 weeks (**Figs. 2F-G**), reflecting early activation followed by late-stage dysfunction.

### Longitudinal shifts in T cell subsets during tumor progression

Cytokines and transcription factors orchestrate T cell polarization over time^35^. For example, at 1 week, *IL-12α, IL-12β* (p < 0.01), and signal transducer and activator of transcription 4 (*STAT4*; p < 0.01) indicated robust type 1 CD4^+^ T helper and type 1 CD8^+^ cytotoxic T cell (Th1/Tc1) differentiation^36^. At 3 weeks, transient but significant (p < 0.05) increases in *IL-21* and *B-cell lymphoma 6* (*BCL6*) suggest CD8^+^ follicular cytotoxic T cell (Tfc)-like response^37^, while modest yet significant (p < 0.05) increases in *STAT3, IL-6*, and *IL-1β* indicate emergence of IL-17/IL-22 producing CD8^+^ Tc17/Tc22-like cells^38-40^ (**Fig. 3A**). By 6 weeks, significant (p < 0.01) induction of *Aryl hydrocarbon receptor* (*AhR*) (p < 0.05) and sustained *IL-1β* expression reinforced Tc17/Tc22 polarization ^41^. In contrast, a significant (p < 0.05) decline of *forkhead box P3* (*FOXP3*) and *STAT5*α at 6 weeks indicates functional loss of Foxp3^+^CD8^+^ Tregs^42^ (**Fig. 3B**).

**Figure 3.**
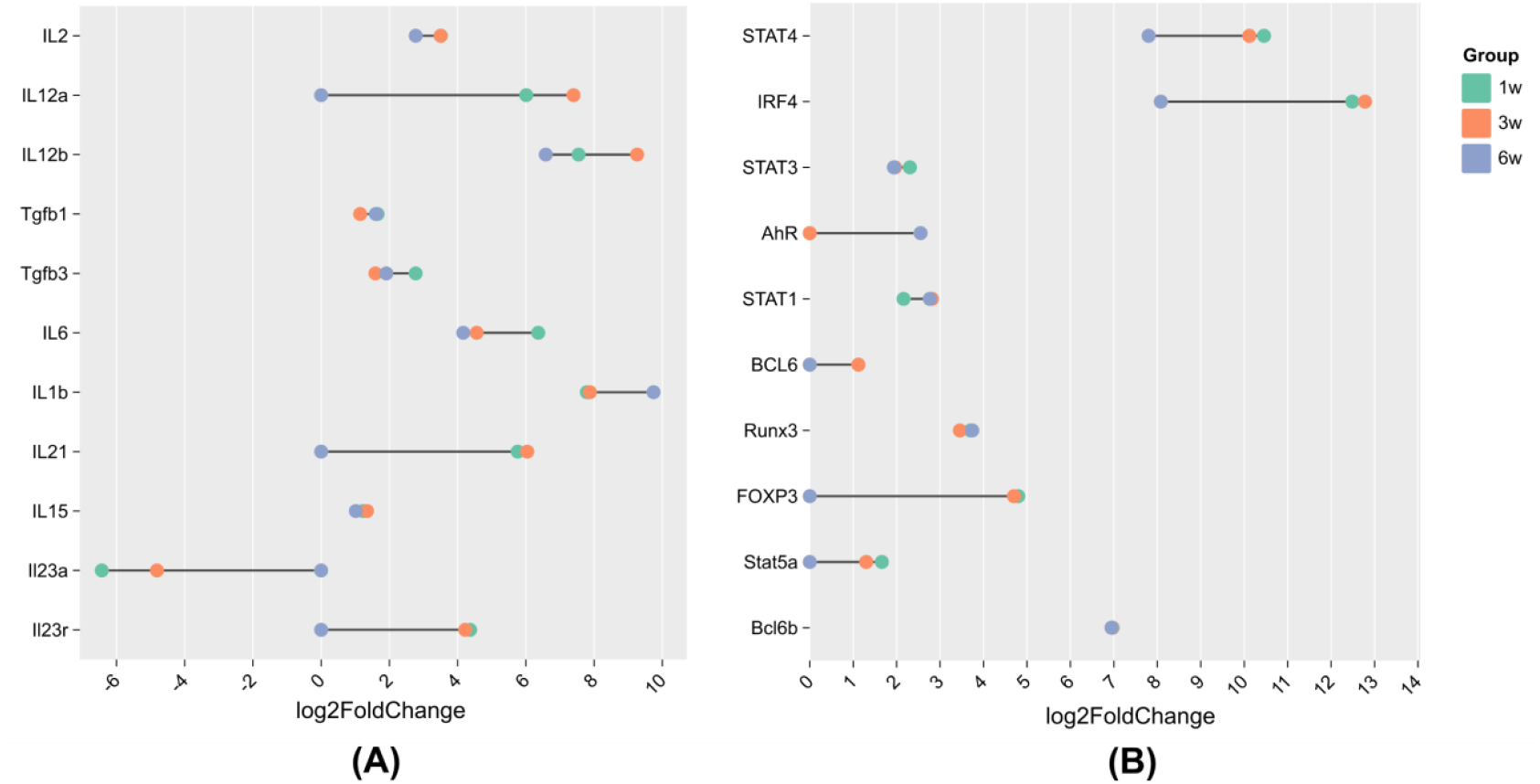
Expression patterns of T cell-related genes associated with (A) polarization and (B) transcription factors. A list of genes associated with T cell polarization and transcription can be found in Koh et al. 2023^43^.

### Immune cytokine and chemokine remodeling underlies T cell plasticity during tumor growth

Pro-inflammatory cytokines were dynamically regulated **(Fig. 4, Table 2**). For instance, IFN-γ, *GZMB* and *PRF1* expression remained elevated across all time points, suggesting a transient peak of Th1/Tc1 activity ^44,45^. Th2 cytokines *IL-5* and *IL-13* were downregulated at 6 weeks (*IL-5*: 5.88 to 0; *IL-13*: 10.69 to 5.67; p<0.01 vs. 1 week). IL-21, a cytokine implicated in CD8^+^ Tfc-like responses, peaked at 3 weeks but was absent at 6 weeks, suggesting transient CD8^+^ Tfc activation that is often associated with B cell help and early tumor immune surveillance ^46^. Parallel to pro-inflammatory cytokines, the immunosuppressive cytokine-related genes underwent substantial remodeling. For example, *IL-10* (6.24 to 3.53) and *indoleamine 2*,*3-dioxygenase 1* (*IDO1*; 6.78 to 4.53) decreased over time, whereas *arginase-1* (*ARG1*) remained high (9.14 to 9.61) **(Fig. 4, Table 2)**.

**Table 1.**
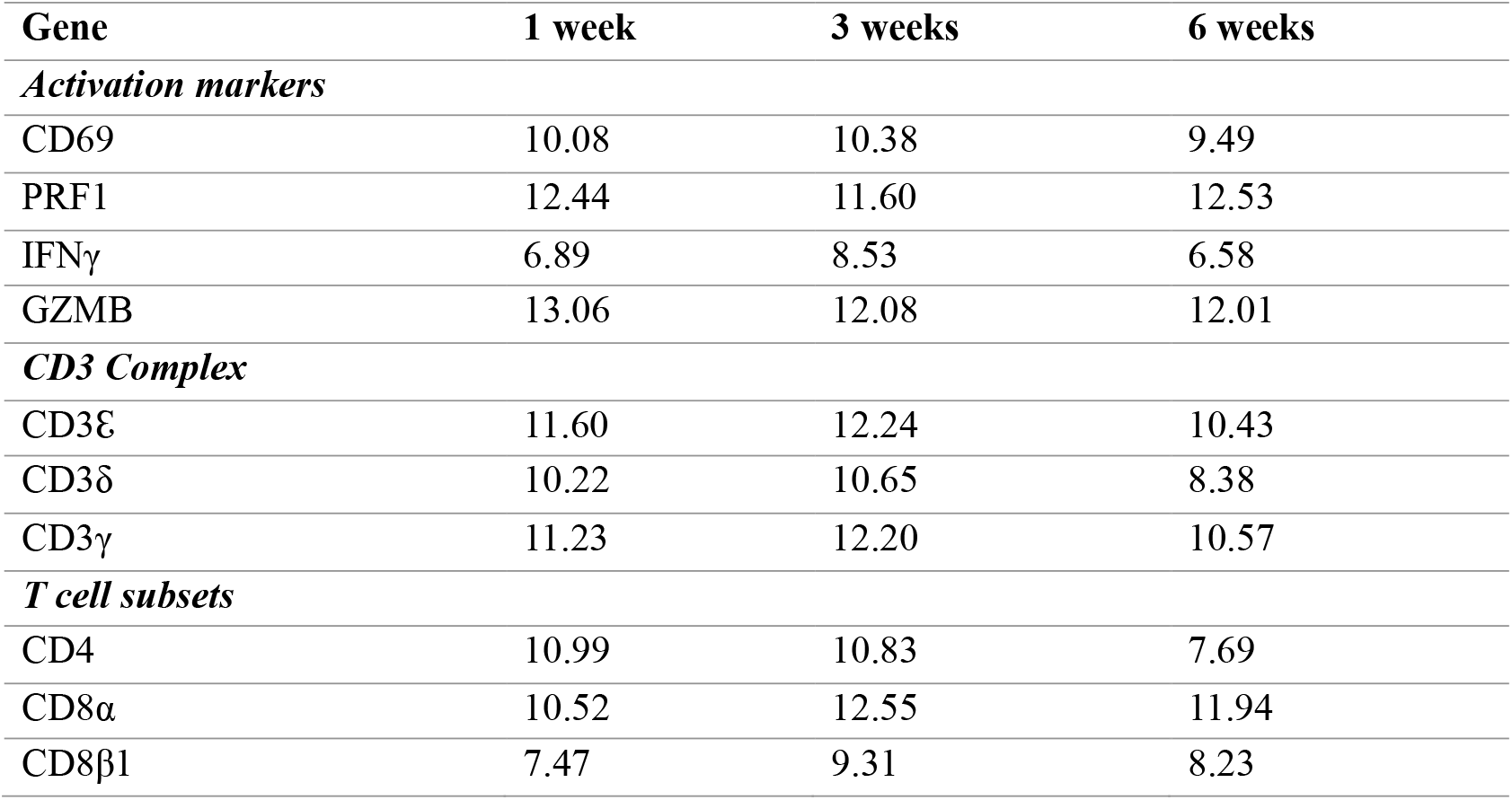
Log2FC of T cell lineage and their activation markers across different stages of tumor growth.

**Table 2.** Log2FC of T cell-associated proinflammatory and immunosuppressive cytokines and chemoattractants.

| Gene | 1 week | 3 weeks | 6 weeks |
| --- | --- | --- | --- |
| ***Proinflammatory cytokines*** |  |  |  |
| IFNG | 6.888 | 8.529 | 6.583 |
| PRF1 | 12.442 | 11.602 | 12.526 |
| IL5 | 4.741 | 5.882 | 0 |
| IL13 | 9.767 | 10.685 | 5.674 |
| Il9r | 6.35 | 7.02 | 3.538 |
| Il21 | 5.76 | 6.04 | 0 |
| IL10 | 6.243 | 5.879 | 3.529 |
| TGFβ1 | 1.65 | 1.143 | 1.6038 |
| TGFβ3 | 2.772 | 1.599 | 1.91 |
| GZMB | 13.06238 | 12.07848 | 12.01156 |
| ***Immunosuppressive cytokines*** |  |  |  |
| IL10 | 6.243 | 5.879 | 3.529 |
| TGFβ1 | 1.65 | 1.143 | 1.6038 |
| TGFβ3 | 2.772 | 1.599 | 1.91 |
| IDO1 | 5.273 | 6.78 | 4.527 |
| ARG1 | 9.406 | 9.14 | 9.611 |
| ***Chemoattractants*** |  |  |  |
| CXCL9 | 10.221 | 11.218 | 10.551 |
| CXCL10 | 1.9208 | 2.4938 | 3.171 |
| CCL17 | 7.738 | 7.904 | 3.395 |
| CCL22 | 13.453 | 13.02 | 9.693 |

**Figure 4.**
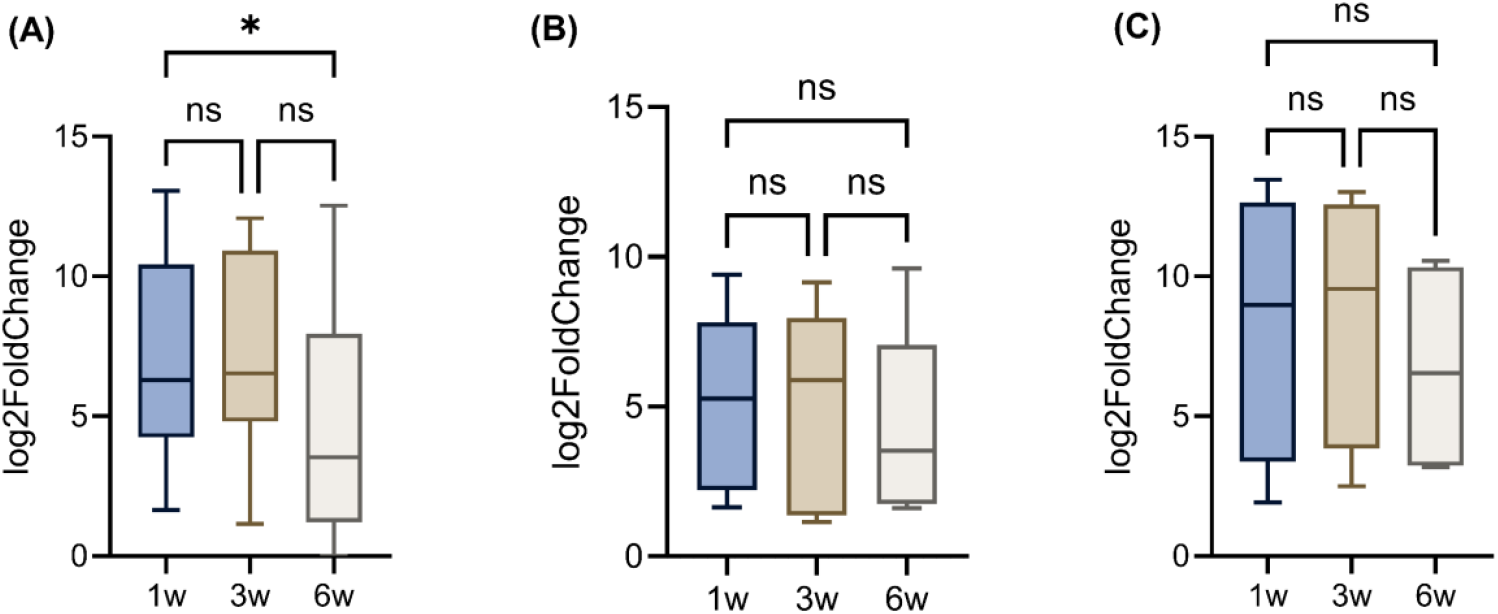
Log2FC of genes related to (A) pro-inflammatory cytokines, (B) immunosuppressive cytokines, and (C) chemoattractants. ns, not significant; w, weeks; *p ≤ 0.05, **p ≤ 0.01, ***p ≤ 0.001, and ****p ≤ 0.0001. Genes presented in these figures are exactly refelcted in **table 2**.

Chemokine profiling offers additional insight into the temporal dynamics of immune cell trafficking. For instance, *C-X-C motif chemokine ligand 9* (*CXCL9*), a potent chemoattractant for CXCR3^+^CD8^+^ T cells and CD4^+^ Th1 cells^47^, remained consistently elevated over time (10.22 to 11.22; p<0.05), suggesting sustained recruitment of cytotoxic effectors. *CXCL10* levels increased linearly over time (p < 0.05), peaking at 6 weeks (3.17), which reinforces the notion of increasing effector T cell infiltration during tumor progression (**Table 2**). Conversely, Treg-attracting regulatory chemokines, such as *CCL17* and *CCL22*, were significantly downregulated over time (*CCL17*: 7.90 to 3.40; *CCL22*: 13.45 to 9.69; p < 0.01). Interestingly, group-wise comparisons revealed that only pro-inflammatory cytokines were significantly upregulated at 6 weeks (p<0.05). In contrast, changes in immunosuppressive cytokines and chemoattractants were not significant (**Figs. 4A-C**).

### Stable inhibitory checkpoint expression with progressive loss of co-stimulation

Recruited T cells within the TME often exhibit features of functional exhaustion due to chronic antigen stimulation and persistent exposure to immunosuppressive cues. This exhausted phenotype is typically marked by sustained expression of multiple inhibitory immune checkpoints, such as programmed cell death 1 (PD-1), T-cell immunoglobulin and mucin domain-containing molecule 3 (TIM-3), lymphocyte activation 3 (LAG-3), and cytotoxic T lymphocyte antigen 4 (CTLA-4), that collectively dampen T cell effector functions and promote immune tolerance^48^. Recent evidence, however, highlights the phenotypic plasticity of these exhausted T cells, indicating that exhaustion exists on a continuum rather than as a terminal state^49^. In this study, exhaustion markers *PD-1, CTLA-4, LAG3, T-cell immunoreceptor with immunoglobulin and ITIM domains* (*TIGIT*), and *B and T lymphocyte attenuator* (*BTLA*) remained stable over 6 weeks (p = 0.24) **(Figure 5A)**. In contrast, co-stimulatory molecules *CD28* and *ICOS* were significantly downregulated at 6 weeks compared to the 1- and 3-week time points (p<0.01 and p<0.001, respectively) **(Fig. 5B, Table 3)**, indicating progressive functional compromise of TILs. **(Table 3)**. These findings support a model in which TILs become increasingly entrenched in a functionally compromised state, marked by stable expression of inhibitory receptors and diminishing co-stimulatory signaling, which are hallmarks of terminal exhaustion^50,51^.

**Table 3.** Log2FC of T-cell exhaustion (and co-stimulation) markers.

| Gene | 1 week | 3 weeks | 6 weeks |
| --- | --- | --- | --- |
| *Exhaustion Markers* |  |  |  |
| PD-1 | 9.277 | 10.89 | 10.99 |
| CTLA-4 | 11.032 | 10.743 | 7.949 |
| LAG3 | 6.358 | 6.103 | 4.229 |
| TIGIT | -1.116 | 0 | 0 |
| BTLA | 12.314 | 11.637 | 9.657 |
| *Co-stimulatory markers* |  |  |  |
| CD28 | 5.482 | 5.738 | 3.168 |
| ICOS | 12.371 | 12.255 | 9.948 |
| CD27 | 4.153 | 4.265 | 1.944 |

**Figure 5:**
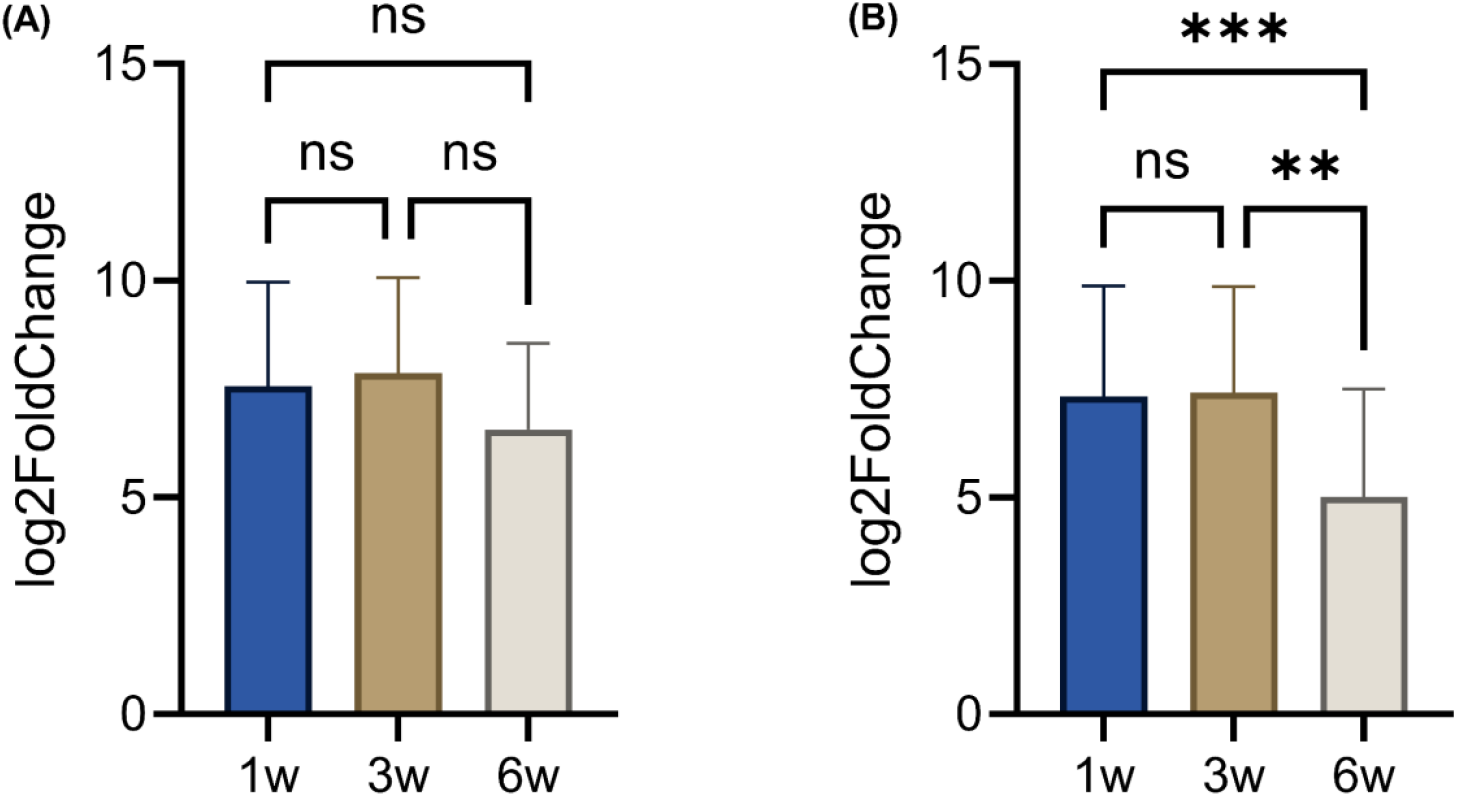
Log2FC of T cell-related genes for (A) exhaustion and (B) co-stimulatory markers. ns, not significant; w, weeks; *p ≤ 0.05, **p ≤ 0.01, ***p ≤ 0.001, and ****p ≤ 0.0001. Genes presented in these figures are exactly refelcted in **table 3**.

### Crosstalk between T cells and APCs

Antigen-presenting cells (APCs), such as dendritic cells (DCs), serve as the primary sentinels of the immune system by capturing debris from dying tumor cells, processing these antigens, and presenting them on MHC class I or II molecules to activate CD8^+^ cytotoxic or CD4^+^ helper T cells, respectively^52^. The functionality of early-stage APCs was evident from high expression of C-C chemokine receptor type 7 (*CCR7*; 7.347), which is essential for APC migration to draining lymph nodes^53^, and MHC Class II genes H2-Aa (15.485) and H2-Ab1 (10.636) at 1 week. By 6 weeks, the expression of these molecules, including Cd1d1/2, ICAM1, H2-Ob, and CD74, declined, indicating a loss of classical antigen-presenting function (**Fig. 6A, Table S5**). Simultaneously, the reduced expression of pattern recognition receptors (PRRs), such as nucleotide-binding oligomerization domain 1 (NOD1) and nucleotide-binding oligomerization domain 2 (NOD2), suggests diminished innate immune sensing and inflammatory priming. These reductions became significant at 6 weeks (p < 0.001), while changes between 1 and 3 weeks were not significant, indicating a delayed but coordinated suppression of APC function **(Fig. 6A)**.

**Figure 6.**
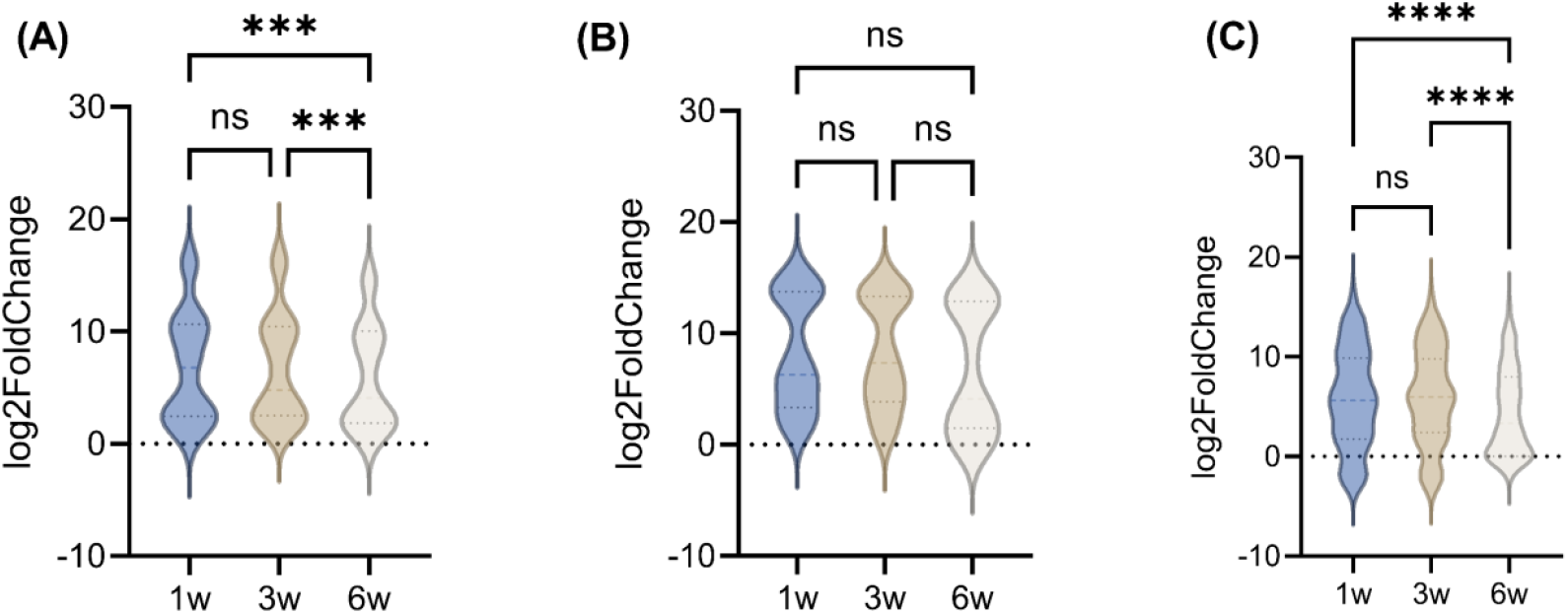
Log2FC of (A) antigen processing cell (APC) genes, (B) dendritic cell genes, and (C) B cell genes. ns, not significant; w, weeks; *p ≤ 0.05, **p ≤ 0.01, ***p ≤ 0.001, and ****p ≤ 0.0001. Genes presented in these figures are exactly refelcted in **Supplement tables S5-S6**.

Elevated expression of *CD83, CD86, CD40, CCR2, and CCL19* at 1 week indicates a mature, pro-inflammatory DC phenotype actively engaged in antigen presentation and migration. High levels of *CCR2, CCR5*, and *CXCR4* at 1 week further supported active DC trafficking toward lymphoid tissues and tumor sites, while increased *IL-10* may reflect early regulatory feedback within an immunostimulatory context **(Table S6)**. These transcriptional patterns suggest robust DC-T cell crosstalk at 1 week. However, as the tumor progresses from 3 to 6 weeks, the DC-T cell axis deteriorates sharply. By 6 weeks, critical mediators such as *CCL19, CCL5*, and *CD40L* were lost, although groupwise comparisons between 1 and 6 weeks did not reach statistical significance (p = 0.0724) **(Fig. 6B, Table S6)**. The loss of these mediators from 3 to 6 weeks suggests a functional collapse in DC-mediated immune activation in late-stage tumors. This plasticity likely reflects a tumor-driven shift toward immune evasion, which is increasingly recognized as a facilitator of metastatic progression^54-56^.

This progressive decline in DC function also impacts the broader adaptive immune network, particularly B cells and macrophages, which rely on effective antigen presentation and cytokine cues from DCs and T helper cells. Mature DCs play a pivotal role in shaping B cell responses by presenting unprocessed antigens in lymphoid follicles, producing supportive cytokines (e.g., IL-6, IL-12), and licensing Tfh cell differentiation via co-stimulatory signals, such as CD40-CD40L^57,58^. Although gene expression associated with B cell function remained relatively stable between 1 and 3 weeks (p = 0.37), a marked and significant downregulation occurred by 6 weeks compared to both earlier time points (p < 0.0001) **(Fig. 6C, Table S7)**. These changes highlight a functional collapse of B cell-mediated immunity as the TME transitions to a more immunosuppressive state, consistent with tumor-driven immune escape mechanisms.

In the immunologically dynamic context of TNBC, macrophages (also recognize as APCs) represent a central hub of immune regulation, toggling between pro-inflammatory (M1-like; anti-tumoral) and immunosuppressive (M2-like; pro-tumoral) states^59,60^. Classical M1-associated genes such as *IL-1β* (7.79), *CCL5* (2.57), and *CXCL9* (10.22) were strongly expressed and remained elevated or slightly increased at 3 and 6 weeks (**Fig. 7A**). Chemokine receptors essential for monocyte and T cell recruitment, such as *CCR2* (15.16), *CCR5* (13.85), and *CCR7* (7.35), were also upregulated early, indicating an active immune infiltration landscape. Transcriptional regulators such as *STAT1* (2.16 to 2.81) and *toll-like receptor 1* (*TLR1*) remained consistently expressed, pointing to stable innate immune activation.

**Figure 7.**
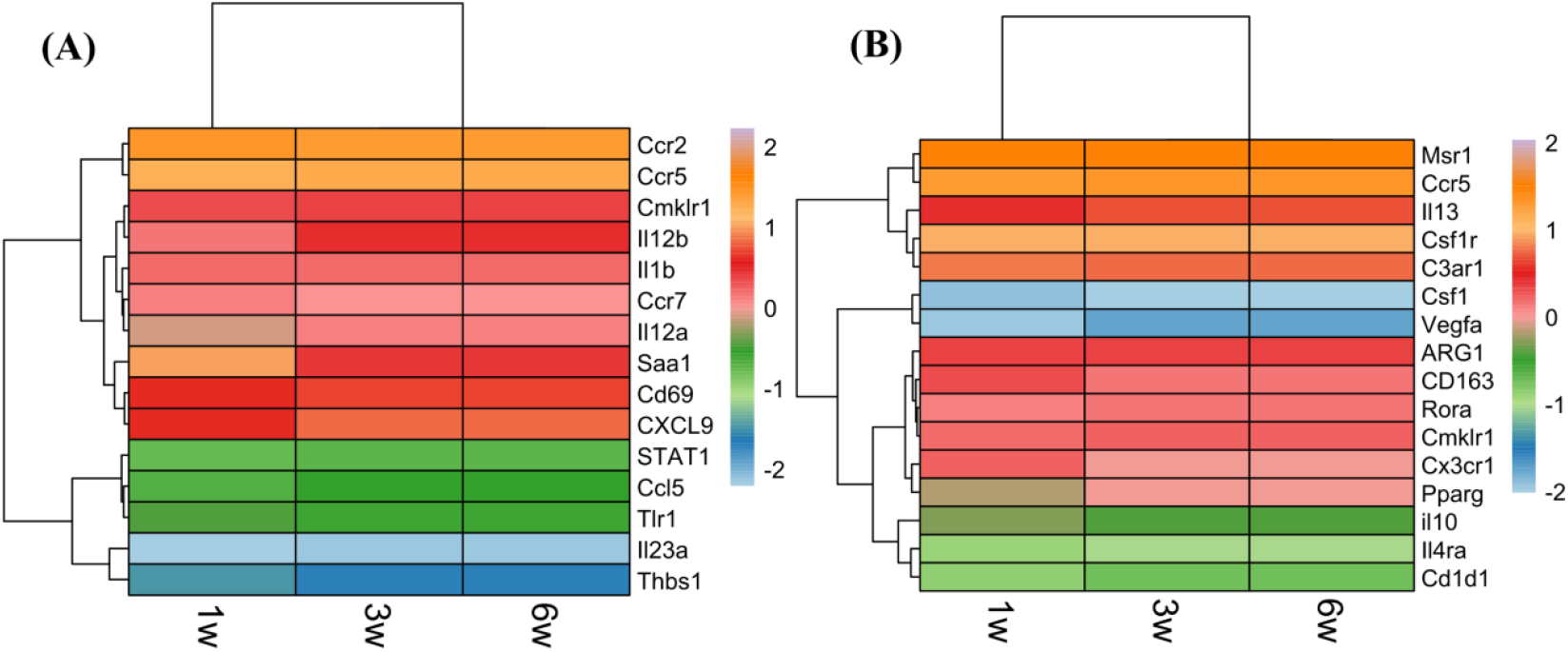
Heatmap of macrophage-associated genes during 4T1 tumor growth. **(A)** M1-associated genes, and **(B)** M2-associated genes; where with orange to red shades indicating upregulation and green to blue shades indicating downregulation. Gene expression levels are shown at the indicated time points of tumor progression, with the scale bar representing normalized values from –2 (lowest expression) to +2 (highest expression).

In contrast, markers associated with M2 polarization followed a distinct temporal trajectory. Colony stimulating factor 1 (*CSF1*), a key M2-polarizing cytokine^61^, remained significantly downregulated across all time points (p < 0.0001), while its receptor, colony-stimulating factor 1 receptor (*CSF1R*), remained highly expressed (p < 0.0001), suggesting an environment that favors macrophage persistence but limits complete M2 differentiation. The M2 cytokine *IL-13* increased at the 3-week time point compared to the 1-week time point (9.77 to 10.69), suggestive of transient Th2-mediated skewing^62^ (p < 0.01). Co-receptors, such as *IL-4Rα* (3.47 to 3.11) and chemokine-like receptor 1 (*CMKLR1*; 8.60 to 8.69), remained stably expressed (p < 0.05), indicating persistent, but not escalating M2 signaling potential. Notably, Arg1, a hallmark of metabolic immunosuppression, remained highly expressed across the time course (p < 0.0001), implying continuous arginine depletion and T cell suppression^63,64^. Additional M2 markers, including *CD163, C3AR1*, and *RORα*, also exhibited sustained expression (p < 0.001), reinforcing the presence of tolerogenic macrophages ^65,66^. Interestingly, *IL-10* expression declined from 6.24 at earlier stages to 3.53 at late stages (p < 0.0001), suggesting reduced immunosuppressive cytokine signaling in advanced tumors (**Fig. 7B**).

## Discussion

Our study demonstrates that T cell plasticity within the 4T1 TNBC TME undergoes profound remodeling during tumor progression, with implications for both tumor adaptation and therapeutic response. Early stages of tumor growth were marked by robust T cell activation, including expression of cytotoxic mediators such as PRF1, GZMB, and IFN-γ, and dynamic polarization supported by IL-12 and STAT4. However, by 6 weeks post-implantation, we observed a collapse in transcriptional breadth across T cell subsets, narrowing of TCR diversity, and downregulation of NKT and γδ TCR genes, consistent with clonal restriction and functional decline (**Fig. 2, Tables S2-S3**). These temporal changes suggest that the tumors progressively erode T cell functionality, shifting the immune balance from early surveillance toward late-stage dysfunction.

While exhaustion markers, such as PD-1 and CTLA-4, remained stably expressed across tumor progression, co-stimulatory molecules, including CD28 and ICOS, were significantly downregulated by 6 weeks, highlighting a state of persistent checkpoint signaling coupled with a loss of co-stimulation (**Fig. 5, Table 3**). This reflects a transition toward terminal exhaustion, which limits the capacity of TILs to respond to antigen stimulation. Concurrently, shifts in cytokine and chemokine expression revealed transient peaks in Th1/Tc1 and Tfc-like activity at early and intermediate stages, followed by suppression of IL-21, IL-13, and IL-15, alongside sustained ARG1 and TGF-β signaling (**Figs. 3-4, Table 2**). These findings illustrate the plasticity of T cell subsets and the reprogramming of immune cytokine networks that underlie the progression from immune activation to immune suppression.

Cross-talk with APCs further reinforced this immune decline. At 1 week, DCs displayed mature, immunostimulatory phenotypes, characterized by high expression of CCR7, H2-Aa, H2-Ab1, CD83, CD86, and CD40, supporting productive T cell priming (**Fig. 6A, Tables S5-S6**). By 6 weeks, however, classical antigen presentation genes, chemokines (e.g., CCL19 and CCL5), and co-stimulatory mediators, including CD40L, were markedly reduced, indicating a functional collapse of DC-T cell interactions (**Fig. 6B, Table S6**). This deterioration extended to B cells, which showed significant downregulation of activation and antigen-presenting pathways by 6 weeks, reflecting impaired humoral immunity (**Fig. 6C, Table S7**). Macrophages maintained a mixed phenotype, e.g., M1-like cytokines (CXCL9, CCL5) persisted, M2-associated markers (ARG1, CD163) were sustained, and IL-13 was transiently upregulated (**Fig. 7**). These findings highlight the emergence of a hybrid macrophage phenotype, maintaining pro-inflammatory features but progressively tolerogenic, which contributes to immune escape and tumor progression.

Together, these observations emphasize that T cell plasticity within the TNBC TME is not static but evolves, shifting from initial activation toward progressive exhaustion, impaired antigen presentation, and immune suppression. This dynamic immune adaptation provides tumors with a dual advantage – evading early cytotoxic responses while creating conditions favorable for metastatic progression. Importantly, our results suggest that the timing of therapeutic interventions is critical. Strategies that restore co-stimulatory signaling, counteract ARG1-mediated metabolic suppression, or sustain DC function may prove most effective when deployed before terminal exhaustion dominates.

This study has limitations, as transcriptomic profiling alone cannot fully elucidate the epigenetic and metabolic mechanisms underlying T cell plasticity. Whole-genome sequencing, single-cell transcriptomics, proteomics and integrative multi-omics approaches will be crucial for further elucidating how T cells and APCs are reprogrammed within the TME. Additionally, while the 4T1 murine model captures many features of human TNBC, future validation in patient-derived models will be needed to establish translational relevance.

## Conclusion

we provide a comprehensive analysis of T cell molecular plasticity during untreated 4T1 TNBC progression, revealing a trajectory from early immune activation to late-stage exhaustion and immune collapse. By defining the transcriptional programs that shape T cell dysfunction and APC decline, this study highlights key vulnerabilities that could be exploited for therapeutic reprogramming. Targeting the timing and mechanisms of plasticity may be central to improving immunotherapy outcomes in TNBC.

## Acknowledgments

D.S. was supported in part by a fund from the Division of Academic Affairs at North Carolina A&T State University, while R.H.N. was supported in part by a grant from the NIH (1R35GM153737).

## Author Contributions

M.I.: Performed data analysis, prepared figures and tables, and wrote the manuscript; R.H.N.: Molecular biology and cell signaling; S.H.H. and R.B.J.: Bioinformatics and data modeling; P.M.M.: Breast cancer genomics; B.L.H.: Preclinical cancer immunology; M.T.H.: Overall guidance to the first author; C.J.R.: 4T1 model system; M.D.T.: Evolutionary genomics; J.L.G.: Evolutionary biology and genomics; H.L.K.: Clinical cancer immunology; D.S.: Conceptualization, study outline, supervision, and data analysis. All authors edited the manuscript and approved the submission.

## Data Availability

Data comprising sequence read counts and differentially expressed genes (DEGs) for the 1-, 3-, and 6-week periods are deposited under DOI: 10.5281/zenodo.16886936.

