## Supplement File for "Molecular Plasticity of T Cells Informs Their Possible Adaptation in 4T1 Tumors"

**Supplementary Tables**

**Table S1:** NCBI SRA run accession numbers of 4T1 TNBC cells and tumors at different growth stages (1-, 3-, and 6-week).

| **Tumor** | **4T1 (1w growth)** | **4T1 (3w growth)** | **4T1 (6w growth)** | **4T1 cells (Baseline)** |
| --- | --- | --- | --- | --- |
| T1 | SRR10428182 | SRR10428188 | SRR10428194 | SRR12898539 |
|  | SRR10428183 | SRR10428189 | SRR10428195 | SRR12898540 |
| T2 | SRR10428184 | SRR10428190 | SRR10428196 | SRR21229974 |
|  | SRR10428185 | SRR10428191 | SRR10428197 | SRR21229975 |
| T3 | SRR10428186 | SRR10428192 | SRR10428198 | ERR3825656 |
|  | SRR10428187 | SRR10428193 | SRR10428199 | ERR3825657 |

**Table S2:** Log2FC of various natural killer T cell (NKT) associated genes at multiple stages of tumor growth (1-, 3-, and 6-week).

| **Gene** | **1w** | **3w** | **6w** |
| --- | --- | --- | --- |
| CD28 | 5.48282 | 5.738377 | 3.168785 |
| PDCD1 | 9.277767 | 10.89942 | 10.99552 |
| CTLA4 | 11.0323 | 10.74373 | 7.949671 |
| ICOS | 12.37176 | 12.25559 | 9.94823 |
| SH2D1A | 9.825447 | 9.124122 | 6.972575 |
| FYN | 0 | 1.010119 | 0 |
| S1PR1 | 5.751489 | 6.00132 | 4.846163 |
| CD1D1 | 3.6 | 4.5 | 1.517 |
| PTPRC | 13.71 | 13.276 | 12.44 |
| CD1D2 | 6.797 | 4.783 | 4.069 |
| IL13 | 9.767 | 10.685 | 5.674 |
| RORA | 8.22 | 8.33 | 6 |
| TBX21 | 8.084 | 8.22 | 5.983 |
| ITGAL | 10.875 | 10.53 | 10.51 |

**Table S3:** Log2FC of various TCR-related genes at different stages of tumor growth (1-, 3-, and 6-week).

| **Gene** | **1w** | **3w** | **6w** |
| --- | --- | --- | --- |
| TRAV1 | 0 | 0 | 5.593635 |
| TRAC | 5.215333 | 5.818741 | 4.191889 |
| TRBV1 | 8.223628 | 9.013937 | 6.66843 |
| TRBV2 | 7.620352 | 7.52806 | 0 |
| TRBV12-2 | 6.314885 | 6.7541 | 5.220269 |
| TRBV14 | 6.654585 | 7.753989 | 5.442985 |
| TRBV16 | 5.627443 | 5.26385 | 0 |
| TRBV17 | 6.012832 | 7.150547 | 0 |
| TRBV19 | 7.793487 | 8.555032 | 7.745628 |
| TRBD1 | 5.573766 | 6.158652 | 0 |
| TRBJ1-1 | 7.956201 | 7.291704 | 0 |
| TRBJ1-2 | 7.296597 | 6.890175 | 0 |
| TRBJ1-3 | 6.587233 | 6.403852 | 0 |
| TRBJ1-4 | 6.159444 | 5.705379 | 0 |
| TRBJ1-5 | 6.509109 | 5.72096 | 0 |
| TRBJ1-6 | 6.728048 | 6.080104 | 5.923745 |
| TRBJ2-1 | 5.889026 | 5.50952 | 0 |
| TRBC1 | 9.755333 | 9.811743 | 8.556481 |
| TRBC2 | 11.81991 | 12.26641 | 10.58005 |
| TRBV1 | 8.223628 | 9.013937 | 6.66843 |
| TRBV13-1 | 7.224374 | 8.146406 | 7.649576 |
| TRBV14 | 6.654585 | 7.753989 | 5.442985 |
| TRBV19 | 7.793487 | 8.555032 | 7.745628 |
| TRBV29 | 6.657896 | 5.876541 | 5.4798 |
| TRBJ1-6 | 6.728048 | 6.080104 | 5.923745 |
| TRBC1 | 9.755333 | 9.811743 | 8.556481 |
| TRBC2 | 11.81991 | 12.26641 | 10.58005 |
| TRBV12-2 | 6.314885 | 6.7541 | 5.220269 |
| TRAV1 | 0 | 0 | 5.593635 |
| TRAV3-3 | 0 | 5.916922 | 6.470688 |
| TRDC | 10.48543 | 9.922193 | 6.755593 |
| CD247 | 6.011 | 5.78 | 3.379 |
| ITK | 8.28 | 8.93 | 6.93 |
| CARD11 | 9.88 | 9.13 | 8.06 |

**Table S4:** Log2FC of various gamma delta T cell-related genes.

| **Gene** | **1w** | **3w** | **6w** |
| --- | --- | --- | --- |
| TRGV1 | 6.56639 | 6.498448 | 0 |
| TRGV2 | 7.08854 | 8.576397 | 9.305937 |
| TRGV6 | 5.633385 | 0 | 0 |
| TRDV4 | 4.974927 | 5.395416 | 0 |
| TRDV5 | 4.620844 | 0 | 0 |
| CD3δ | 10.22224 | 10.65418 | 8.375881 |
| CD3ℇ | 11.60537 | 12.24258 | 10.4261 |
| CD3γ | 11.23106 | 12.1983 | 10.57474 |
| CD247 | 6.011193 | 5.780067 | 3.379392 |
| CD28 | 5.48282 | 5.738377 | 3.168785 |
| CD27 | 4.153751 | 4.265074 | 1.944618 |
| IL2 | 3.509571 | 3.501903 | 2.772738 |
| IL7 | -1.82706 | -1.57025 | -1.3076 |
| IL15 | 1.231285 | 1.345062 | 1.016437 |
| IL22 | 13.63109 | 13.23528 | 13.78705 |
| IFNγ | 6.888718 | 8.529825 | 6.583763 |
| SOX13 | 3.205401 | 3.208396 | 0 |
| ZBTB16 | 10.46515 | 12.14555 | 6.135456 |
| TBX21 | 8.08405 | 8.228109 | 5.983699 |
| BCL11B | 5.994845 | 6.895263 | 4.068734 |
| LCK | 2.354114 | 2.472996 | 0 |
| FYN | 0 | 1.010119 | 0 |
| ZAP70 | 9.622031 | 9.543039 | 7.767536 |
| LAT | 5.94311 | 6.321327 | 4.080068 |
| LCP2 | 13.99676 | 13.90644 | 13.33247 |
| ITK | 8.289076 | 8.937925 | 6.934933 |
| NFATC1 | 0 | 0 | 1.130898 |
| CCR6 | 5.991487 | 5.726745 | 0 |
| CCR9 | 6.189079 | 5.830836 | 0 |
| ITGAE | 4.309458 | 3.83807 | 2.72139 |
| S1PR1 | 5.751489 | 6.00132 | 4.846163 |
| TRDC | 10.48543 | 9.922193 | 6.755593 |
| TRGC1 | 8.827772 | 9.105033 | 7.354446 |
| TRGC2 | 8.077999 | 9.301093 | 9.945164 |
| CD4 | 10.9952 | 10.83271 | 7.693341 |
| CD8α | 10.51535 | 12.55518 | 11.94493 |
| CCR5 | 13.84574 | 13.42237 | 13.41786 |
| CXCR4 | 13.25985 | 12.82128 | 12.4328 |
| ICOS | 12.37176 | 12.25559 | 9.94823 |
| KLRK1 | 11.60687 | 11.47893 | 10.43512 |
| FAS | 1.561955 | 1.235113 | 1.210129 |
| GzmA | 14.87896 | 12.29685 | 9.082225 |
| GzmB | 13.06238 | 12.07848 | 12.01156 |
| GzmC | 7.726491 | 7.871223 | 13.15444 |
| PRF1 | 12.44229 | 11.60221 | 12.52697 |
| TNF | 3.639242 | 4.307926 | 2.172809 |
| IL23R | 4.367571 | 4.226369 | 0 |
| IL18R1 | 10.00721 | 9.630116 | 8.070308 |
| CD83 | 14.03598 | 14.12054 | 12.21179 |
| CLEC7A | 16.95106 | 15.88612 | 15.00325 |

**Table S5:** Log2FC of various antigen-presenting cell (APC)-related genes.

| **Gene** | **1w** | **3w** | **6w** |
| --- | --- | --- | --- |
| CCR7 | 7.347 | 6.88 | 4.23 |
| CD1D1 | 3.605 | 4.5 | 1.517 |
| CD1D2 | 6.797 | 4.783 | 4.069 |
| CD74 | 7.657 | 7.812 | 6.273 |
| FCER1g | 10.977 | 10.435 | 10.09 |
| FCGR1 | 9.55 | 9.411 | 10.05 |
| FCGR2b | 16.296 | 15.92 | 14.96 |
| FCGR3 | 11.2869 | 10.854 | 10.327 |
| H2-Aa | 15.485 | 15.813 | 13.767 |
| H2-Ab1 | 10.636 | 10.696 | 8.912 |
| H2-D1 | 1.912 | 2.124 | 2.026 |
| H2-DMa | 4.29 | 4.214 | 3.368 |
| H2-DMb1 | 11.212 | 10.98 | 10.613 |
| H2-DMb2 | 10.157 | 9.787 | 9.202 |
| H2-Eb1 | 16.38 | 16.748 | 14.764 |
| H2-K1 | 1.535 | 1.44 | 1.535 |
| H2-Ob | 10.4 | 10.308 | 8.08 |
| H2-Q1 | 0 | 1.55 | 1.846 |
| H2-Q2 | 1.538 | 1.324 | 1.833 |
| H2-T23 | 2.64 | 2.716 | 0 |
| ICAM1 | 3.308 | 3.049 | 0 |
| NOD1 | 2.878 | 3.11 | 1.972 |
| NOD2 | 2.441 | 2.006 | 2.055 |
| PSMB9 | 2.565 | 2.624 | 2.638 |
| SLC11A1 | 8.8 | 8.438 | 7.799 |
| TAP1 | 2.42 | 2.52 | 2.644 |
| TAPBP | 1.216 | 1.294 | 1.328 |

**Table S6:** Log2FC of various dendritic cell (DC)-related genes.

| **Gene** | **1w** | **3w** | **6w** |
| --- | --- | --- | --- |
| CCL19 | 5.889 | 7.365 | 0 |
| CCL5 | 2.568 | 3.846 | 0 |
| CCR1 | 13.63 | 13.235 | 13.78 |
| CCR2 | 15.157 | 14.258 | 13.211 |
| CCR5 | 13.84 | 13.422 | 13.417 |
| CD40 | 3.337 | 3.35 | 2.84 |
| CD40L | 8.595 | 8.954 | 0 |
| CD83 | 14.035 | 14.12 | 12.211 |
| CD86 | 13.76 | 13.32 | 12.89 |
| CXCR1 | 6.278 | 6.155 | 8.45 |
| CXCR4 | 13.2598 | 12.821 | 12.432 |
| IL10 | 6.243 | 5.879 | 3.529 |
| LYN | 1.628 | 1.281 | 1.498 |
| TGFβ1 | 1.65 | 1.143 | 1.603 |
| CD80 | 4.657 | 4.548 | 4.128 |

**Table S7:** Log2FC of various B cell-related genes.

| **Gene** | **1w** | **3w** | **6w** |
| --- | --- | --- | --- |
| ATM | -1.882 | -1.609 | -1.165 |
| BLK | 6.367 | 6.525 | 4.013 |
| BLNK | 8.014 | 7.474 | 5.792 |
| BMI1 | -1.789 | -1.583 | 0 |
| BTLA | 12.314 | 11.637 | 9.657 |
| CARD11 | 9.88 | 9.131 | 8.069 |
| CASP3 | -2.3 | -2.45 | 0 |
| CD19 | 5.301 | 8.263 | 6.348 |
| CD200R1 | 9.931 | 8.953 | 7.201 |
| CD22 | 11.483 | 11.409 | 9.27 |
| CD27 | 4.153 | 4.265 | 1.944 |
| CD28 | 5.482 | 5.738 | 3.168 |
| CD37 | 4.795 | 4.613 | 2.36 |
| CD38 | 0 | 0 | 2.062 |
| CD4 | 10.995 | 10.832 | 7.693 |
| CD40 | 3.337 | 3.352 | 2.843 |
| CD40L | 8.595 | 8.954 | 0 |
| CD69 | 10.078 | 10.38 | 9.486 |
| CD70 | 0 | 6.414 | 0 |
| CD74 | 7.657 | 7.812 | 6.273 |
| CD79α | 4.044 | 4.425 | 1.466 |
| CD79β | 6.755 | 6.858 | 4.836 |
| CD83 | 14.035 | 14.12 | 12.21 |
| CD86 | 13.762 | 13.328 | 12.896 |
| CTLA4 | 11.032 | 10.743 | 7.949 |
| CXCL13 | 9.689 | 11.867 | 10.632 |
| CXCR5 | 3.075 | 4.192 | 2.203 |
| DPP4 | 11.45 | 11.754 | 8.381 |
| FAS | 1.561 | 1.235 | 1.21 |
| FCGR2B | 16.296 | 15.929 | 14.964 |
| FLT3 | 7.738 | 6.741 | 4.019 |
| FOXP3 | 4.802 | 4.706 | 0 |
| GPR183 | 7.11 | 6.417 | 6.041 |
| ICOSL | 2.337 | 2.302 | 1.543 |
| IKZF1 | 12.464 | 12.177 | 11.165 |
| IL10 | 6.243 | 5.879 | 3.529 |
| IL11 | -2.905 | -2.92 | 0 |
| IL13 | 9.767 | 10.685 | 5.674 |
| IL1R2 | 5.368 | 6.418 | 6.832 |
| IL21 | 5.76 | 6.04 | 0 |
| IL2Rα | 12.186 | 11.975 | 9.672 |
| IL2Rγ | 3.559 | 3.35 | 0 |
| IL5 | 4.741 | 5.882 | 0 |
| IL6 | 6.362 | 4.558 | 4.164 |
| IL7 | -1.827 | -1.57 | -1.307 |
| IL7R | 13.98 | 13.451 | 11.635 |
| IL21 | 5.76 | 6.04 | 0 |
| INPP5D | 6.172 | 5.899 | 5.692 |
| JAK3 | 1.608 | 1.67 | 2.727 |
| LCK | 2.354 | 2.472 | 0 |
| LYN | 1.628 | 1.281 | 1.498 |
| MEF2C | 0 | 2.525 | 0 |
| MIF | -2.483 | -2.876 | 0 |
| MS4A1 | 7.476 | 9.306 | 6.876 |
| NT5E | -2.148 | -2.133 | 2.342 |
| PRDM1 | 9.133 | 8.965 | 8.862 |
| PRKCD | 1.771 | 1.375 | 0 |
| PTPRC | 13.713 | 13.276 | 12.444 |
| SH2B2 | 1.189 | 1.789 | 0 |
| SYK | 13.548 | 13.297 | 11.908 |
| TGFB1 | 1.65 | 1.143 | 1.603 |
| TIRAP | -1.313 | -1.293 | 0 |
| TNFRSF13B | 10.2 | 9.586 | 8.854 |
| TNFRSF13C | 2.598 | 2.519 | 0 |
| TNFRSF4 | 2.852 | 3.114 | 0 |
| TNFSF13B | 4.705 | 5.725 | 4.77 |
